# Cholesterol Myth and the Imperative Role of Functional Evaluation of Lipoproteins in Atherosclerotic Risk Prediction Strategies

**DOI:** 10.1101/559625

**Authors:** Ravi Divya Bhavani, Srinivasan Ashokkumara, Velusamy Prema, S Narasimhan Kishore Kumara, Chakrapani Lakshmi Narasimhan, Abhilasha Singh, Mohan Thangarajeswari, Kannan Thiruvengadam, Moongilpatti Angappa Arumugam, Periandavan Kalaiselvi

**Author notes:** **Corresponding author: Dr. P. Kalaiselvi** Associate Professor, Department of Medical Biochemistry, Dr ALM Post Graduate Institute for Basic Medical Sciences, University of Madras, Chennai, India. Mail ID.

## Abstract

**Objective:** To arrive at apposite risk prediction strategies in identifying markers that could delineate patients susceptible for atherosclerotic progression from other diabetic patient population thus circumventing undesirable drug exposure or conversely inadequate treatment.

**Research Design and Methods:** The study was designed to assess and compare the biochemical parameters along with lipoprotein ratios assessed in structural and functional aspects between the probable risk factor groups for CAD(diabetics with no symptoms for CAD) and untreated CAD patients (on admission for CAD) in a total of 593 participants. Anthropometric measurements were made and serum fasting glucose, lipid peroxide levels, lipid and lipoprotein ratio assessments were carried out. In addition to these basic parameters, oxidized LDL (Ox-LDL) and serum paraoxonase (PON) activity were also measured in this study. To ascertain the diagnostic accuracy of the individual markers, receiver operating characteristic curve analysis was carried out.

**Results:** Serum levels of blood lipids, lipid peroxides were higher in the diabetic and the cardiovascular patients with a concomitant decrease in the levels of HDL. Increased median levels of Ox-LDL and decreased PON activity was observed in the cardiovascular patients both with and without diabetes.

**Conclusion:** This study proposes Ox-LDL/PON assessment would be more functional than the traditional LDL-c/HDL-c to categorize the risk group of patients into prospective atherosclerotic predisposed and non-predisposed population, thus enabling the former to get treated with appropriate medication.

## Introduction

Diabetes mellitus is regarded to be acquiring swiftly the status of being a most prospective threat to the human race globally, as the morbidity and mortality levels due to its latent complications pose significant health management hardships on our kith and kin. Research in diabetes constantly draws attention not only because the preponderance of diabetes is constantly on the rise and about a twofold increase is estimated worldwide by 2030 (Wild et al., 2004, Whiting et al., 2011) but also due to the impact it confers on the socio-economic status of the society involved (Yesudian et al., 2014).

Diabetes mellitus can be described as a metabolic disease of diverse aetiology characterized by chronic hyperglycemia often found to co-exist with biomolecular disarray as a consequence of either or both impaired insulin secretion and action. Hyperglycemia generates reactive oxygen species (ROS) causing damage to the cells which is considered as one of the most imperative mechanism leading to biomolecular injure and dysregulation at the cellular level (Chang and Chuang, 2010). Furthermore oxidative stress has a profound versatile role in sustaining the metabolic and molecular abnormalities leading to other complications (Giacco and Brownlee, 2010). In particular, the increased glycosylation of proteins like low-density lipoprotein (LDL) has been shown to augment its susceptibility to oxidation and elicit the atherosclerotic processes (Mohan and co-workers, 2010). The modus operandi behind this elevated risk may be in part accredited to the discrepancy between free radicals and antioxidants causing biomolecular stress and damage. Even though diabetes is well attributed to house the conditions of oxidative stress along with lipid derangements and that various epidemiological data have provided plausible evidence that the risk of CAD is increased by the presence of diabetes, (Chait and Bornfeldt, 2009) it is undeniable that not all the diabetics succumb to atherosclerosis. Quite a few studies divulge that diabetic status may perhaps not be a CAD equivalent in all circumstances, thus implicating the requirement of multivariate analysis as an appropriate method for risk stratification in the prevention of cardiovascular complications in patients with diabetes (Bertoluci and Rocha, 2017). Lipid profile analysis along with the evaluation of the lipoprotein ratios is regarded to be the customary laboratory investigation employed to assess the atherogenic risk intensity and to determine the medication strategy that usually aims at lowering the LDL cholesterol / increasing the HDL cholesterol (Hadjiphilippou and Ray, 2018). Although this assessment of lipid and lipoproteins is quite less cumbersome and commonly employed, the biomarker potentials of various lipoproteins for the risk stratification remain contentious (Upadhyay, 2015).

Various indices of the oxidative stress predictors like Oxidised LDL (Ox-LDL), the modified form of LDL which is atherogenic in nature that try to demarcate the diabetic subjects prone to atherosclerotic events from those who do not do so have been analyzed by researchers but disease prediction has been unpredictable because of the conventional approach of considering a single biomarker rather than considering multiple biomarkers and their interactions. Even though several studies validate Paraoxonase (PON), a functional protein of HDL to reflect the oxidative status of the body, (Hofer et al, 2006) it has failed to emerge as a routine analytical parameter with diagnostic significance due to lack of combinatorial assessment. Not only evolution of new biomarkers with improved prediction accuracy would be a mostvalued contribution towards diagnostic assessments, but also precise risk prediction in disease prevention and management at a cost effective manner with simple biochemical assays may have a societal impact on the diagnosis of disease burden.

Biomarkers like Ox-LDL and PON on which adequate research has been performed to designate their role in atherogenesis have not been simultaneously assessed in the risk and disease states. In this regard, a cross sectional case-control study was designed to co-evaluate the functional assessment of lipoproteins as Ox-LDL /PON, the pivotal modulators involved in atherogenic progression in comparison with the routine traditional markers like LDL and HDL and furthermore statistically emerge with a better predictor of risk. Presently there exists no definite reference interval for Ox-LDL levels and the PON enzyme activity. Hence this study was premeditated in diabetics and coronary artery disease (CAD) patients to appraise the roles of individual factors contributing to atherosclerosis and identify the most relevant assay that delineates them to high and low risk.

## Methods

Ethical clearance for the study was obtained from the Institutional Human Ethics Committee and those who consented to take part in the study were recruited from The Government Kilpauk Medical College and Hospital, Chennai from the year 2015-2017 (IHEC Approval No: UM/IHEC/08-2013-I). Patients diagnosed of CAD and are admitted in the hospital with c tnT/ c tnI levels >0.01 ng/ml; patients with the duration of diabetes ≥4years were recruited whereas, patients with other cardiac disorders, HIV infection and other chronic diseases, like tuberculosis, thyroid problems, liver & kidney disorders were excluded from the study. The healthy subjects with no history of cardiovascular disease or diabetes mellitus were included as control population in the study.

Anthropometric measurements of height, weight were taken and body mass index (BMI) was calculated. Venous blood samples were drawn at baseline from the subjects under sterile condition after overnight fasting of 8-12 hrs. Serum/Plasma were separated and stored at −80°C for subsequent measurements. Routine biochemical investigations were carried out using standard biochemical kits for ascertaining the normal functioning of the kidney and liver. The concentration of plasma Ox-LDL was determined with cusabio competitive ELISA kit according to manufacturer’s instructions and the concentration was read at 450 nm. Serum PON activity was assessed by measuring the capacity of the enzyme to liberate p-nitro phenol based on the absorbance recorded at 412 nm during the enzymatic reaction kinetic. Blanks devoid of enzyme were used to exact for the spontaneous hydrolysis of the substrate.

### Statistical Analysis

All statistical analyses were performed using SPSS (Statistical Package for the Social Sciences) software version 20. Values were presented as n (%) and median (Range). Assessment using the – Shapiro-Wilk W test showed that the data was not normally distributed. Comparisons between groups were done using the Non-Parametric Kruskal-Wallis followed by Dunn’s post-hoc test (Table 1) and significance was determined at the level of 5%.The values of PON and Ox-LDL among the groups were compared by area under the receiver operating characteristic (ROC) curves (Figure 1). The Youden Index was estimated to find the cut point with the maximum of sensitivity and specificity. Sensitivity was plotted on the Y axis and the proportion of false positives (1-specificity) on the X axis. Optimal cut-offs were derived from the ROC curves by maximizing the sum of sensitivity and specificity. ROC curves are measures used to estimate the diagnostic accuracy of the individual markers. Area under the curve (AUC) is considered to be the value of accuracy for diagnosing a disease. It is considered to be more predictive if it is closer to 1 while the parameter is regarded to lose its diagnostic significance when it is less than 0.5.

**Table 1:** Represents the Baseline Characteristics and the Biochemical Parameters of the Study Population. PON-Paraoxonase; Ox-LDL-Oxidized LDL; LDL-Low-density lipoprotein; HDL-High density lipoprotein; TCHOL-Total Cholesterol; AIP (Atherogenic Index of Plasma) – Log (TG/HDL-c).

| **Factors** | **Control**<br>**(n=111)** | **DM**<br>**(n=230)** | **CAD + DM**<br>**(n=122)** | **CAD**<br>**(n=130)** | **Sig.** |
| --- | --- | --- | --- | --- | --- |
| **Age** | 56 (40 - 70) | 56 (41 - 70) | 56 (40 - 69) | 56 (40 - 70) | 0.813 |
| **Gender** |  |  |  |  |  |
| **Male** | 31 (27.9%) | 62 (27%) | 31 (25.4%) | 31 (23.8%) | 0.884 |
| **Female** | 80 (72.1%) | 168 (73%) | 91 (74.6%) | 99 (76.2%) |  |
| **Height** | 1.63 (1.44 - 1.88) | 1.63 (1.46 - 1.89) | 1.62 (1.45 - 1.89) | 1.64 (1.42 - 1.86) | 0.161 |
| **Weight** | 72 (45 - 111) | 70 (48 - 111) | 70 (50 - 109) | 70.5 (50 - 111) | 0.575 |
| **BMI** | 26.75 (20.66 - 35.22) | 26.67 (20.17 - 36.58) | 26.64 (20.6 - 35.14) | 26.45 (20.31 - 35.22) | 0.625 |
| **Glucose** | 89 (62 - 142) | 152 (78 - 688) | 167.5 (100 - 210) | 90 (58 - 136) | <0.001 |
| **TChol** | 164 (121 - 180) | 215 (123 - 288) | 250.5 (159 - 298) | 235.5 (164 - 275) | <0.001 |
| **TGL** | 122 (96 - 140) | 168 (100 - 266) | 206 (104 - 278) | 188 (104 - 235) | <0.001 |
| **HDL** | 58 (30 - 77) | 36 (20 - 64) | 24.57 (19.41 - 32) | 30 (19 - 53) | <0.001 |
| **VLDL** | 24.4 (19.2 - 28) | 33.5 (3.4 - 53.2) | 41.2 (20.8 - 55.6) | 37.6 (20.8 - 47) | <0.001 |
| **LDL** | 83.6 (33 - 117.2) | 144.88 (57.4 - 233.8) | 184.5 (107.2 - 228.65) | 170.01 (108.4 - 210.8) | <0.001 |
| **LPO** | 3.1 (1.1 - 4.7) | 12 (2.5 - 26.5) | 26 (16 - 36) | 21 (17 - 25) | <0.001 |
| **PON** | 530 (289 - 585) | 384.69 (295 - 532) | 204 (170 - 416) | 247 (231 - 487) | <0.001 |
| **OXLDL** | 2.11 (1 - 16.5) | 29.45 (2.6 - 63.1) | 74.2 (22.8 - 80) | 58.95 (25.1 - 72.8) | <0.001 |

**Figure 1:**
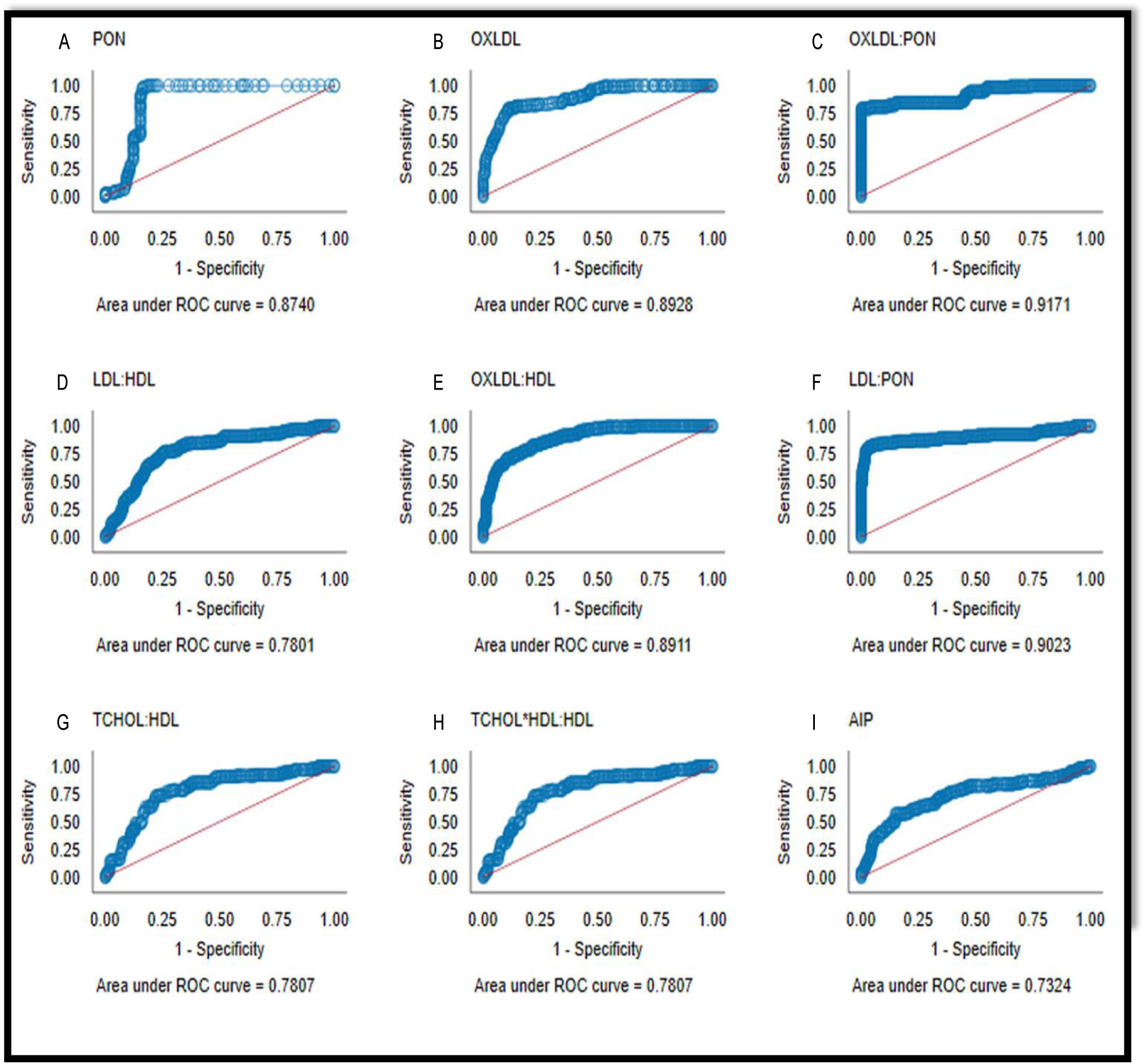
Receiver-operating characteristic (ROC) curves for the biomarkers. **A** – ROC curve of PON among the study population; **B** – ROC curve of Ox-LDL among the study population; C – ROC curve of PON: Ox-LDL among the study population; The AUC values was appreciative for PON and Ox-LDL both independently as well in ratio combination (AUC values-0.87; 0.89; 0.91-respectively); **D** – ROC curve of LDL: HDL among the study population. Although traditional methods substantiate the evaluation of LDL and HDL cholesterol levels for atherogenic risk assessment, the sensitivity and specificity was comparatively low for the LDL: HDL ratio; **E** – ROC curve of OX-LDL: HDL among the study population; **F** – ROC curve of LDL: PON among the study population. When the functional aspects of the lipoproteins were substituted for their levels, the resulting ratios had a significant change in the predictive values with their AUC values being 0.89 and 0.90 respectively; **G** – ROC curve of TCHOL: HDL among the study population; **H** – ROC curve of (TCHOL-HDL): HDL among the study population; **I** – ROC curve of AIP among the study population. For comparative reasons, ROC curve analysis was performed for other ratios commonly used in atherogenic risk stratifications. The AUC values of these ratios were found to be low (AUC values-0.78; 0.78; 0.73-respectively) with a marked loss in sensitivity.

## Results and Discussion

Baseline characteristics and the biochemical parameters of the entire sample population are shown in Table 1. The mean age and BMI at baseline was found to be 56.0 years (range, 40 to 70) and 26 (range, 20.17 − 36.58) respectively. All participants included in this study harmonized for age, gender percentage and BMI. In accordance with the WHO guidelines the median total cholesterol and LDL was found to be optimal in the control individuals while the levels of TC, TGL, VLDL and LDL among the diabetic and CAD patients were found to be increased. However HDL cholesterol was found to be low in patients with diabetic and cardiovascular conditions.

In the present study, marked elevation in the fasting blood glucose levels signifies the status of poor glycemic control being prevalent amongst the diabetic patients involved. Generally in diabetes there exists anomalous metabolism of triglyceride-rich lipoproteins associated with increased production or decreased clearance of VLDL particles which is considered decisive in the pathophysiology of atherogenic dyslipidemia (Krauss, 2004). Similarly our findings are also in line with the conviction made from several other studies that the abnormalities in lipid and lipoproteins are quite renowned in diabetes mellitus as well as CAD and thus may perhaps have a role to play in causing atherosclerosis through more than one route (Renard et al., 2004).This noxious combination of hyperlipidemic status is often found to co-exist with yet another deadly element, the oxidative stress leading to vascular complications in the pathophysiology of diabetes (Pham-Huy et al., 2008).Our results also showed increased levels of lipid peroxides in the diabetic and cardiovascular patients. Evidences censure that the ROS generation and alterations in redox signaling to be the tactical mediator of vascular inflammation in atherogenesis (Pitocco et al., 2013).The LDL are more susceptible and are major targets for oxidative modification and furthermore oxidatively modified form of LDL (Ox-LDL) play imperative role in atherogenic disease progression. Hence further investigations were directed towards Ox-LDL.

Serum Ox-LDL levels were considerably increased in the cardiovascular patients than in control and diabetic population. The median Ox-LDL level in the study population was found to be about fourteen fold increased in the diabetics whereas in cardiovascular patients there was about twenty nine fold while, the diabetic patients with CAD had about thirty seven fold increase when compared to that of the control patients. Our result was in agreement with the literature which suggests the imperative role of oxidative modification of LDL in atherogenic disease progression particularly in conditions of abnormal lipid levels and disturbed antioxidative capacity (Toth, 2014; Alique et al., 2015). On the other hand, the diabetic population without CAD complications had only 1.7 fold increase in the LDL levels while CAD with or without diabetes presented a 2.0-2.2 fold increase in the LDL levels when compared to the controls substantiating the large differences in the Ox-LDL levels compared to that of LDL, which is considered to be one of the prognostic indicator for progression of atherosclerosis.

Also the activity of PON enzyme whose levels reflect the oxidative status was found to be markedly diminished in the cardiac and diabetic patients. Similar kind of observations denoting a diminution in its activity or level to be associated with increased oxidative stress in serum was shown by Kontush and Chapman (2010) and Brites et al., (2017). The reduction of PON activity was more in the diabetic patients with cardiovascular complications compared to the normal as well as diabetic population.

While the Serum PON enzyme activity was found to be significantly decreased in cardiovascular and the diabetic patients with the median being 204 IU/L and 247 IU/L respectively whereas the median being twofold high (530 IU/L) in normal individuals. On assaying HDL as HDL-cholesterol, the diabetic population presented with 38% lower levels of HDL-cholesterol when compared to control while, the levels were about 59% reduced in the DM+CAD group. Conversely, PON levels were drastically reduced in the CAD patients with or without diabetes while, diabetics without CAD had considerably higher PON activity suggesting that PON rather than HDL cholesterol can be a better predictor. This again validates that assessing HDL in terms of cholesterol alone will not dictate the efficiency or functionality of HDL-Cholesterol. This is supported by the fact that the lipid-lowering therapy that aims to increase HDL cholesterol like CETP inhibitors and niacin failed to show promising results questioning whether HDL-C–raising drugs should be a part of preventive armamentarium. Nevertheless, HDL as a therapeutic target cannot be questioned as they correlate with reduction in CAD risk. This accentuates the fact that the guardian angel conjecture of HDL cholesterol does not absolutely rely on their levels but more specifically on their functional feature which also applies for risk validation and a necessity to assess dysfunctional HDL (Linton et al., 2015).

Although there was significant statistical difference among the groups for parameters like lipid profile, PON and Ox-LDL, in order to single out which ones best envisage disease outcomes and to stratify the risk of cardiovascular complications in the diabetic group, ROC curve analysis was performed. As lipoprotein ratios are commonly employed tool for cardiovascular risk prediction, various ratios like LDL/HDL, (Total Cholesterol-HDL)/HDL, calculated atherogenic index of plasma-log (Triglycerides/HDL-c) were derived and subjected to ROC analysis to validate its sensitivity and specificity. The ratios displayed a disappointingly low sensitivity of 41.5%, 43.1%, 43.1% and 45.4% respectively in differentiating atherogenic and diabetic patients. However, when the functional aspect of both these lipoproteins was substituted for Ox-LDL and/PON, like Ox-LDL/HDL and LDL/PON both sensitivity and specificity were appreciable with superior AUC value. The ROC curve analysis performed presented the laboratory cut-point of best predictive result for differentiating CAD patients from Diabetics to be 54.1 mU/ml (sensitivity = 80.8% and specificity = 85.2%) for Ox-LDL and 300 IU/L (sensitivity = 83.1% and specificity = 98.7%) for PON enzyme activity. It is also suggestive that oxidized LDL and PON, when used in combination as a ratio (PON: Ox-LDL) serves to be a superior biomarker displaying 80.8% sensitivity and 91.3% specificity with the AUC being 0.91 than even the traditional lipoprotein ratio (LDL/HDL) employed in the recent times for discriminating the diabetic patients who are at high risk environment of oxidative stress and lipid derangements and with intense propensity for developing atherosclerotic complications.

## Conclusion

This study proposes Ox-LDL/PON assessment would be functional to categorize the risk group of patients into prospective atherosclerotic predisposed and non-predisposed population, thus enabling the former to get treated with appropriate medication. Furthermore the study imparts that rather than the conventional practice of arriving at the risk prediction with the lipid profile and lipoprotein ratios alone, incorporating their functional aspect would certainly aid in guiding the medical practitioners in the appropriate treatment and management consequently not exposing the needless patients to a plethora of lipid lowering medications that come with their clandestine side-effects.

## Limitations of the Study

This outcome was from a cross sectional study and so further cohort studies in the risk factor patients would be more apposite in assessing the effectiveness of this biomarker.

## Acknowledgments

The financial assistance to **Ravi Divya Bhavani** from University Grants Commission – Basic Scientific Research (UGC-BSR) in the form of Senior Research Fellowship (SRF), New Delhi, India is greatly acknowledged.

## Author Contributions

**Ravi Divya Bhavani** – Designed and executed the experimental work; Sample collection, processing, experimentation and data collection; **Srinivasan Ashokkumar & Kannan Thiruvengadam** – Statistical data analysis & interpretation; **Ravi Divya Bhavani, Velusamy Prema & S Narasimhan Kishore Kumar** – Drafted the article/error and grammatical check; **Chakrapani Lakshmi Narasimhan, Abhilasha Singh & Mohan Thangarajeswari** – Sample Collection from different wards (Diabetic and CVD wards); **Moongilpatti Angappa Arumugam** – Clinical assistance for sample access from patients; **Periandavan Kalaiselvi** – Conception of the work and corresponding author.

### Ethical Approval and Consent to Participate

All experiments were performed in accordance with the guidelines approved by the Institutional Human Ethical Committee (UM/IHEC/08-2013-I).

## References

Alique M, Luna C, Carracedo J, Ramírez R. LDL biochemical modifications: a link between atherosclerosis and aging. Food Nutr Res. 2015 Dec 3;59:29240. doi:10.3402/fnr.v59.29240.

Bertoluci MC, Rocha VZ. Cardiovascular risk assessment in patients with diabetes. Diabetol Metab Syndr. 2017 Apr 20;9:25. doi:10.1186/s13098-017-0225-1. eCollection 2017. Review. Erratum in: Diabetol Metab Syndr. 2017 Sep 19;9:70.

Brites F, Martin M, Guillas I, Kontush A. Antioxidative activity of high-density lipoprotein (HDL): Mechanistic insights into potential clinical benefit. BBA Clin. 2017 Aug 19;8:66-77. doi:10.1016/j.bbacli.2017.07.002. eCollection 2017 Dec.

Chait A, Bornfeldt KE. Diabetes and atherosclerosis: is there a role for hyperglycemia?. Journal of lipid research. 2009 Apr 1;50(Supplement):S335-9.

Chang YC, Chuang LM. The role of oxidative stress in the pathogenesis of type 2 diabetes: from molecular mechanism to clinical implication. Am J Transl Res. 2010 Jun 10;2(3):316–31.

Giacco F, Brownlee M. Oxidative stress and diabetic complications. Circ Res. 2010 Oct 29;107(9):1058-70. doi:10.1161/CIRCRESAHA.110.223545.

Hadjiphilippou S, Ray KK. Lipids and Lipoproteins in Risk Prediction. Cardiol Clin. 2018 May;36(2):213-220. doi:10.1016/j.ccl.2017.12.002.

Hofer SE, Bennetts B, Chan AK, Holloway B, Karschimkus C, Jenkins AJ, Silink M, Donaghue KC. Association between PON 1 polymorphisms, PON activity and diabetes complications. J Diabetes Complications. 2006 Sep-Oct;20(5):322–8.

Kontush A, Chapman MJ. Antiatherogenic function of HDL particle subpopulations: focus on antioxidative activities. Curr Opin Lipidol. 2010 Aug;21(4):312-8. doi:10.1097/MOL.0b013e32833bcdc1.

Krauss RM. Lipids and lipoproteins in patients with type 2 diabetes. Diabetes care. 2004 Jun 1;27(6):1496–504.

Linton MF, Yancey PG, Davies SS, Jerome WG, Linton EF, Vickers KC. The role of lipids and lipoproteins in atherosclerosis. 2015.

Mohan V, Venkatraman JV, Pradeepa R. Epidemiology of cardiovascular disease in type 2 diabetes: the Indian scenario. J Diabetes Sci Technol. 2010 Jan 1;4(1):158–70.

Pham-Huy LA, He H, Pham-Huy C. Free radicals, antioxidants in disease and health. International journal of biomedical science: IJBS. 2008 Jun;4(2):89.

Pitocco D, Tesauro M, Alessandro R, Ghirlanda G, Cardillo C. Oxidative stress in diabetes: implications for vascular and other complications. International journal of molecular sciences. 2013 Oct 30;14(11):21525–50.

Renard CB, Kramer F, Johansson F, Lamharzi N, Tannock LR, von Herrath MG, Chait A, Bornfeldt KE. Diabetes and diabetes-associated lipid abnormalities have distinct effects on initiation and progression of atherosclerotic lesions. J Clin Invest. 2004 Sep;114(5):659–68.

Toth PP. Insulin resistance, small LDL particles, and risk for atherosclerotic disease. Curr Vasc Pharmacol. 2014;12(4):653–7.

Upadhyay RK. Emerging risk biomarkers in cardiovascular diseases and disorders. J Lipids. 2015;2015:971453. doi:10.1155/2015/971453. Epub 2015 Apr 8.

Whiting DR, Guariguata L, Weil C, Shaw J. IDF diabetes atlas: global estimates of the prevalence of diabetes for 2011 and 2030. Diabetes Res Clin Pract. 2011 Dec;94(3):311-21. doi:10.1016/j.diabres.2011.10.029.

Wild S, Roglic G, Green A, Sicree R, King H. Global prevalence of diabetes: estimates for the year 2000 and projections for 2030. Diabetes Care. 2004 May;27(5):1047–53.

Yesudian CA, Grepstad M, Visintin E, Ferrario A. The economic burden of diabetes in India: a review of the literature. Global Health. 2014 Dec 2;10:80. doi:10.1186/s12992-014-0080-x.

